# Does Sex-Differential Gene Expression Drive Sex-Differential Selection in Humans?

**DOI:** 10.1101/2024.07.23.604672

**Authors:** Matthew J. Ming, Changde Cheng, Mark Kirkpatrick, Arbel Harpak

## Abstract

Sex differences in human transcriptomes have been argued to drive sex-differential natural selection (SDS). Here, we show that previous evidence supporting this hypothesis has been largely unfounded. We develop a new method to test for a genome-wide relationship between sex differences in expression and selection on expression-influencing alleles (eQTLs). We apply it across 39 human tissues and find no evidence for a general relationship. We offer possible explanations for the lack of evidence, including that it is due in part to eQTL ascertainment bias towards sites under weak selection. We conclude that the drivers of ongoing SDS in humans remain to be identified.

## Introduction

Sex-differences in gene expression have been theorized to be the *result* of long-term sex-differential selection (SDS), where allelic fitness effects differ between males and females^1,2^. When a gene product is only beneficial in one sex, it is expected that expression modifiers will evolve to increase expression for that sex and decrease expression in the other ^3^. The reverse causality has also been proposed: sex differences in gene expression might be the *driver* of SDS on expression modifiers^1,4^.

SDS acting on viability within the current generation can generate between-sex differences in allele frequency^5,6^ at expression modifiers (expression Quantitative Trait Loci, or eQTLs)^3^. Sexually dimorphic traits (diseases being one example with strong repercussions on survival^6,7,8,9^) may be subject to SDS acting through viability. Previous work has proposed and tested theoretical models for relating divergence in eQTL allele frequencies with differential expression between sexes^1,10,11^. Some of the empirical results have, however, been called into question^12,13,14^.

A controversial result by Cheng and Kirkpatrick (2016)^1^ (hereafter “CK16”) is a characteristic pattern relating between-sex *F*_ST_^1,5,15^ (henceforth “*F*_ST_”) to sex differences in gene expression (henceforth “Δ”). They observed high *F*_ST_ values at intermediate values of Δ and near-zero values of *F*_ST_ when a gene is expressed evenly between the sexes (Δ = 0) or is only expressed in one sex (Δ = ±1). They nicknamed this bimodal pattern “Twin Peaks”. The Twin Peaks pattern has been used as a signal to detect SDS in several species^1,4,16,17^.

Here, we revisit the CK16 model and statistical approach. We find that their interpretation was based on previously unappreciated statistical artifacts. We then refine their model and apply it to new, more extensive data on gene expression and allele frequencies. Across 39 human tissues, we find no evidence for a genome-wide relationship between viability SDS and Δ. We discuss how a bias in eQTL discovery towards eQTLs under weaker selection can explain the lack of signal for a relationship between SDS and Δ, and how the drivers of SDS may still be investigated.

## Results

### Twin Peaks is a Statistical Artifact

We begin by revisiting the Twin Peaks pattern with a critical eye towards caveats in its application and interpretation. Most importantly, we reconsider the model that CK16 proposed to explain the pattern. It is based on two key assumptions. First, the relationship between a gene’s expression levels and its effect on fitness in each sex is linear. Second, sexually antagonistic selection is symmetric, meaning selection coefficients are equal in magnitude and opposite in sign between sexes. This yields a quadratic relationship between *F*_ST_ and Δ at a biallelic site affecting expression. At small absolute values of Δ,

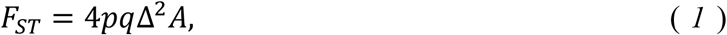

where *p* is the allele frequency in zygotes and *q* = 1 − *p. F*_ST_ is the between-sex fixation index^18^ which is used to quantify allele frequency differences between males and females. In the model, these differences are due solely to the sex differences in post-zygotic fitness effects of alleles. The quantity Δ is the sex difference in gene expression (**Methods**). Finally, *A* is a compound parameter involving the within-sex effect of gene expression level on fitness.

The expectation in CK16 for *F*_ST_ at extreme expression differences (|Δ| → 1), however, is based on intuition rather than a model. The authors suggest that if a gene is not expressed in one sex (Δ = ±1), then selection will not act on it. Selection on the other sex should optimize expression levels, so under the symmetrical selection assumption there will be no ongoing directional selection in that sex either. As neither sex experiences selection, there will be no force driving increased *F*_ST_. CK16 then interpolate a bimodal shape by joining the quadratic relationship at low |Δ| and the expectation for *F*_ST_ = 0 at Δ = ±1 (**Fig. 1**).

**Figure 1:**
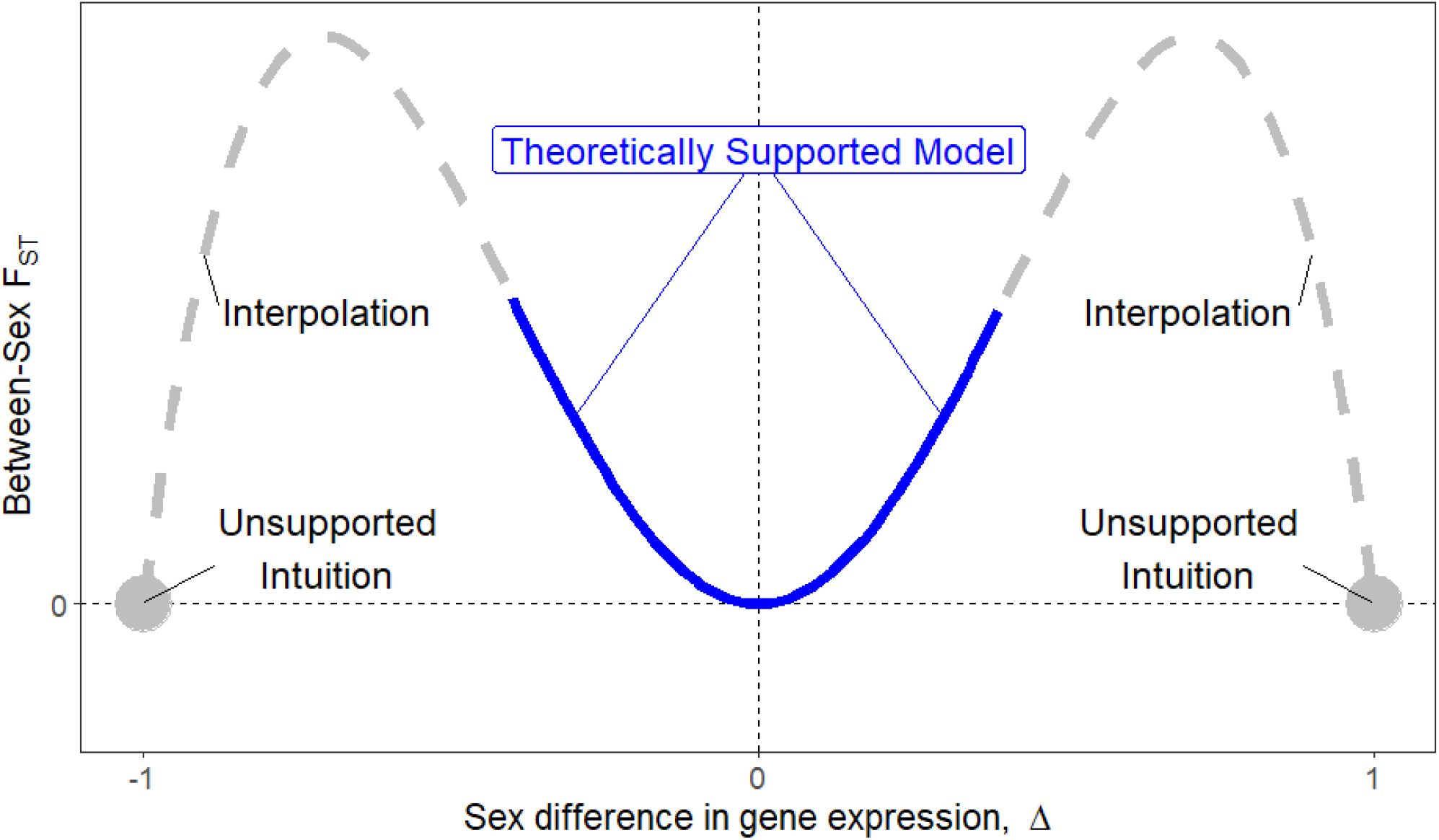
Revisiting the Twin Peaks theoretical expectation. Cheng and Kirkpatrick (2016) derive an expectation for a quadratic relationship at low values of Δ (solid blue line). However, their expectation for “Twin Peaks” is based on a reasoning for F_ST_ = 0 at genes with sex-specific expression (Δ = ±1) that is unsupported. Additionally, values between the theoretically supported region and the extremes are based on qualitative interpolation (dashed line). The portions of the model which are not supported are shown in grey.

However, the assumption of symmetric selection used at low values of Δ may be inappropriate for large Δ values. In particular, when Δ = ±1, the lack of expression in one sex plausibly suggests different selection between males and females. It is therefore not intuitive that *F*_ST_ should simply go to zero at sites regulating expression in these genes. Additionally, although the quantity 2*pq* appears in **Eq. 1**, CK16 did not include that term in fitting the model, effectively assuming it to only contribute random noise to the relationship between *F*_ST_ and Δ.

Because of these caveats to the model, we decided to revisit the support for Twin Peaks. We first ask whether the Twin Peaks pattern is due to SDS by applying the statistical tests of CK16 to data generated under a null hypothesis of no relationship between Δ and SDS. We replicated the pattern shown in CK16 by using the same data and methods. Namely, we performed a 4^th^ degree polynomial regression of *F*_ST_ on Δ, using allele count data from 1000 Genomes^19^ and expression data from the gonads (ovaries and testes) in GTEx v3^20^. The curve and associations generated using these datasets we refer to as the “real data” (blue line in **Fig. 2a**).

**Figure 2.**
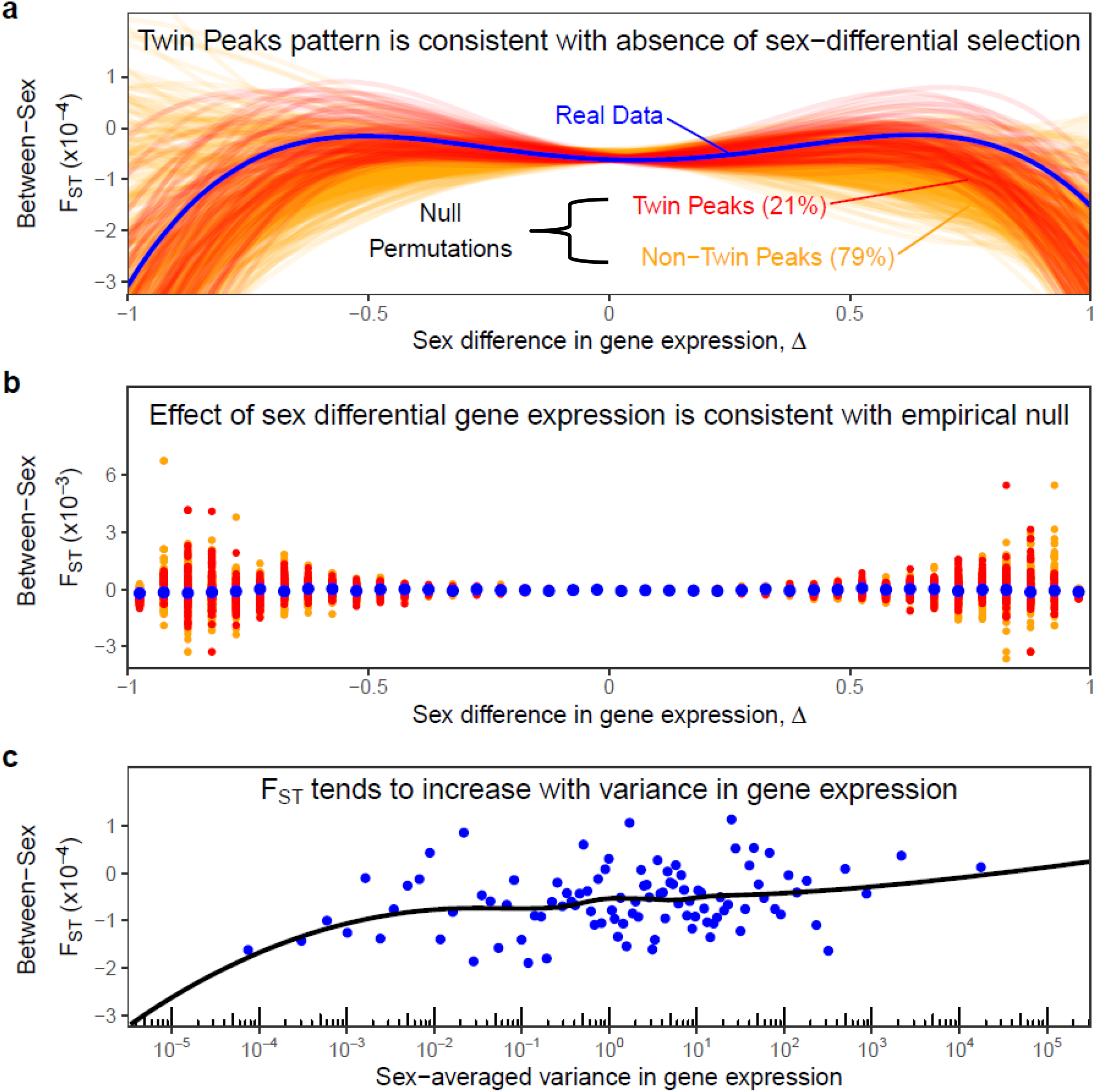
The Twin Peaks pattern is the result of statistical artifacts. **a)** A 4^th^-degree polynomial regression of between-sex *F*_ST_ to sex differences in gene expression for the real data is shown in blue. Red and yellow curves shown are fits based on samples from an empirical null generated by permuting the sex labels in expression data. In red (yellow) are null iterations classified as consistent (inconsistent) with the Twin Peaks expectation according to criteria employed by Cheng and Kirkpatrick (2016). **b)** The data was split into 40 evenly spaced bins of sex difference in gene expression to visualize mean trends. The x-axis values show the midpoint of each bin, and the y-axis values show mean male-female *F*_ST_ of biallelic sites within each bin. **c)** A positive relationship between *F*_ST_ and variance in gene expression (regardless of sex) may contribute to the genome-wide relationship between sex differences in gene expression and selection. Genes are divided into 100 evenly sized bins of sex-averaged sample variance in gene expression. The y-axis values show the mean male-female *F*_ST_ for all biallelic sites for all genes in each the bin. The black curve shows a LOESS fit to the non-binned data.

We then generated an empirical null by permuting ovary and testes tissue labels in the expression data but retaining sex labels associated with *F*_ST_ values (**Methods**), then recomputed Δ values, and again fit a 4^th^-degree polynomial. We find 21% of permutations qualify as Twin Peaks according to the criteria used by CK16, and that the polynomial fit to the real data is not visually distinct from those fits to the null (**Fig. 2a**). Higher order polynomial regressions can also yield spurious fits because distant points have an oversized impact^21,22^. We therefore binned genes by Δ values and examined the relationship with mean *F*_ST_ in each bin. Again, the real data shows no unusual relationship between *F*_ST_ and Δ compared to null data (**Fig. 2b**). From these results, we conclude the Twin Peaks pattern is not statistically significant.

Importantly, this permutation method differs from the method used in CK16, which permuted Δ values across genes. Both methods break the associations between *F*_ST_ and Δ as desired. Our sex -label permutation method, however, preserves *F*_ST_ associations with the gene’s overall expression, maintaining gene features such as expression variance which the CK16 method does not. This explains why their reported *p*-value for Twin Peaks curves (0.016) is lower than our (0.21).

We hypothesized that one reason for the spurious Twin Peaks pattern is confounding. In particular, variance in expression (regardless of sex) is positively correlated with both Δ and *F*_ST_. Previous work has shown that genes subject to weaker stabilizing selection show higher variance in expression^23^. Higher variance means larger differences between randomly selected subgroups, and therefore it should translate to larger values of Δ even in the absence of SDS. In turn, stronger (sex-agnostic) selection can lead to stronger drift at linked sites^24,25,26,27^. Taken together, the confounding with variance in gene expression could generate a relationship between SDS and |Δ| in the absence of SDS. Indeed, expression variance and *F*_ST_ are positively correlated (Pearson *p* = 0.011; **Fig. 2c**), consistent with this hypothesis. In sum, we do not find support for Cheng and Kirkpatrick’s conclusion that there is a genome-wide relationship between *F*_ST_ and Δ.

### No evidence for genome-wide SDS on eQTLs

Although we do not find support for a relationship between sex differences in gene expression and selection using CK16’s methodology for generating Twin Peaks, one may still exist. We test this hypothesis across many tissues using improved statistical modeling, data, and methods. Despite the caveats to portions of the model discussed above, we believe that CK16’s theoretical expectation of a quadratic relationship between *F*_ST_ and small values of Δ is valid. We therefore built on that model by introducing the compound parameter δ^2^ = 4*qp*Δ^2^ and rewriting **Eq. 1** as

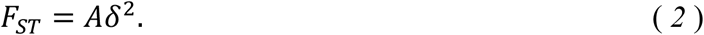

(**Methods**). To estimate *A*, for each gene-tissue pair, we used the single *cis*-eQTL from GTEx v8^28^ with the strongest association with its expression. Our estimator of *A* is then the inverse-variance-weighted linear regression of *F*_ST_ on *δ*^2^ (**Eq. 2**). The advantages of this formulation over CK16’s are that it allows direct estimation of *A*, the strength of SDS on sex differences in expression. This model also accounts for variation in allele frequencies across sites. Further, by using a single eQTL to calculate *F*_ST_ for the whole gene, we circumvent biases which can arise when using a simple mean to estimate gene-wide *F*_ST_^29,30^.

To accompany the updated regression methodology, we also updated the datasets for both allele frequencies and gene expression. For allele frequencies, we used the Non-Finnish European subset in gnomAD v3 (averaging over 60,000 allele samples per site)^31^. We used expression data from 49 tissues from the GTEx v8 dataset^28^ (averaging over 200 male samples and 150 female samples). Both datasets greatly expand our sample size compared to CK16, and GTEx v8 provides sample-specific sex labels for calculating Δ across multiple tissues instead of just the gonads—ovaries and testes—as in the original Twin Peaks paper.

Using the updated statistical framework method, we find no evidence that *A* differs significantly from zero in any tissue (**Fig. 3; Methods**). One explanation for the absence of a pattern may be found in recent work by Mostafavi et al. (2023). The authors contrasted how selection impacts the discovery of genome-wide association study (GWAS) hits with the discovery of eQTLs. Variants with large effects on phenotypes are expected to segregate at low frequencies, reducing discovery power. In GWAS, this is counterbalanced by increased power due to their large effect sizes. Consequently, in GWAS, low frequency variants can still be detected if their effect is large enough. In contrast, eQTL discovery is based only on the effect of genotype on gene expression which does not necessarily translate to fitness-relevant trait variation. Detection of strongly selected sites is therefore less likely^32^. This ascertainment bias can weaken the relationship between Δ and *F*_ST_ at eQTLs, as sites with high *F*_ST_ are less likely to be eQTLs.

**Figure 3.**
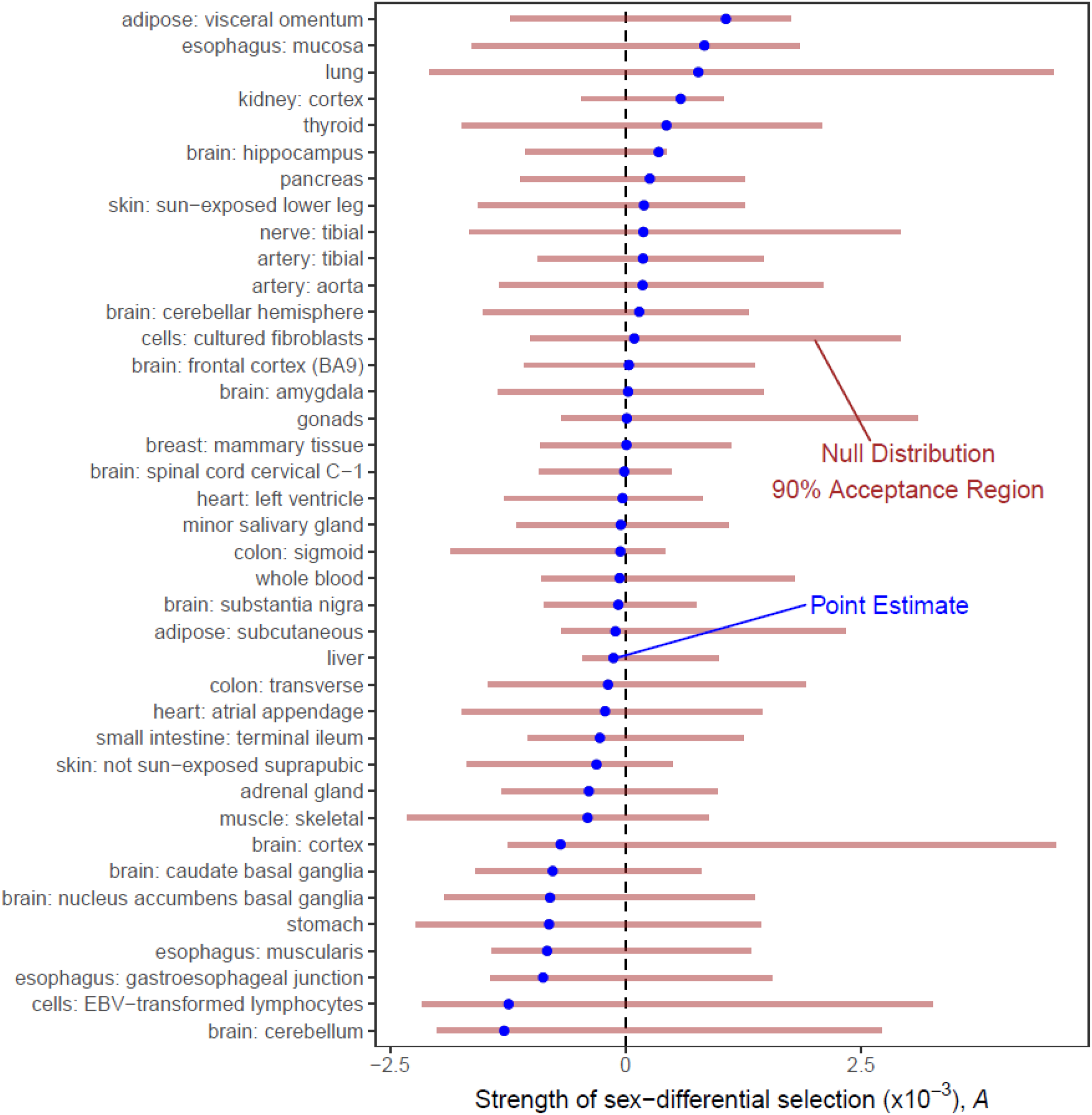
Lack of evidence for sex-differential selection on sex-differential gene expression using newer data. Shown are estimates of *A* (**Eq. 2**). Acceptance regions are based on the 5^th^ to 95^th^ quantiles of an empirical null distribution generated by permuting sex labels in the gene expression data.

In “**Twin Peaks is a Statistical Artifact**”, we suggested that large sex differences in expression may be entirely unrelated to SDS—for instance, merely tagging genes with high expression variance. There are other potential explanations for the lack of a genome-wide relationship. While some sex differences in expression may be driven by or drive SDS, these are the exception rather than the rule. The majority of sex differences in expression may be a regulatory side effect of SDS on different genes. Lastly, sex-differential gene expression may be due to past SDS, but not drivers of current SDS^33^. Regardless of the reason, if a causal relationship between Δ and FST is rare in the genome, it would be difficult to detect using models that assume a pervasive, persistent relationship between the two.

## Conclusion

Previous work suggested a genome-wide relationship between sex differences in expression and SDS. However, this work was based on statistical artifacts and confounded effects, such as stabilizing selection and variance in gene expression. Even when using newer data and improved statistical methods, we found no evidence for a genome-wide relationship between sex-differential expression and contemporary SDS. In contrast, studies that measured SDS irrespective of expression or trait variation have reported pervasive, genome-wide signals of SDS in the human genome^5,10,34^. While causal relationships, past and present, between SDS and sex-differential gene expression in humans remain plausible, they are yet to be fully elucidated.

## Acknowledgements

We thank Jared Cole for comments on the manuscript and other members of the Harpak Lab for helpful conversations. The work was funded by NIH grants GM11685307 to M.K., and R35GM151108 and a Pew Scholarship to A.H.

## Materials and Methods

### Permuting Twin Peaks sex-labels

In the section “**Twin Peaks is a Statistical Artifact**”, we demonstrate that the Twin Peaks curve presented in CK16 is not statistically significant compared to a permuted null. To do so, we used the methods and datasets described in CK16 and **Eq. 1**. We computed allele count using the 1000 Genomes dataset^19^ and calculating *F*_ST_. We filtered out any sites with only a single alternative allele in either males or females (i.e., singletons). We used the Transcripts Per Million (TPM) normalized GTEx v3 (referred to as “pilot” on the download page) dataset^20^ for expression levels for calculating Δ. Because GTEx v3 does not have individual sample labels, this analysis only compared expression in the gonads (ovaries and testes) where expression level summaries represent a single biological sex. We used the Ensembl GRCh38 v77^35^ annotation file for gene annotations. Following CK16, we limited our analysis to protein-coding genes.

To estimate *F*_ST_, we used Hudson’s estimator^36^ based on the R package used in CK16^37^ as presented by Bhatia et al. (2013)^29^,

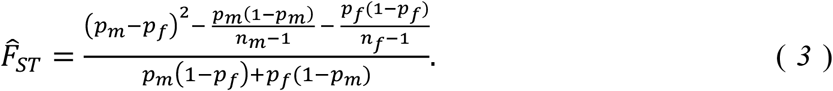

Here, *pm* and *pf* are the allele frequencies in males and females respectively, and *nm* and *nf* are the number of males and females respectively. As a measure of sex-differential expression we used Δ as defined in CK16:

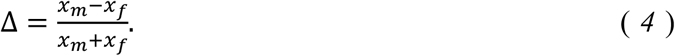

Here, *x*_*m*_ and *x*_*f*_ are sex-averaged TPM-normalized expression levels in males and females respectively. Because 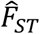 in **Eq. 3** is a site-specific estimator while Δ is gene-wide, we generated a gene-wide estimate of *F*_ST_ by taking a simple average of all sites within a gene body plus 1000bp upstream and downstream. To generate the Twin Peaks curve, we used a 4^th^-degree polynomial regression between 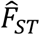 and Δ. We note that here and in CK16, this regression therefore ignores variation in heterozygosity across sites (**Eq. 1**). In our improved model (**Results: “No evidence for SDS on eQTLs across all genes in multiple tissues”** and **Methods: “Applying a new model and method for SDS-expression regression”**) we correct this omission.

To compare the original Twin Peaks curve to curves generated under the null of no SDS, we generated an empirical null distribution of 4^th^-degree regression curves using sex-label permutations of gene expression data. We permuted the ovary and testes labels for each GTEx sample, then recalculated Δ for all genes. We then re-performed the 4^th^-degree polynomial regression on the new Δ values (the *F*_ST_ values remain unchanged). This was repeated 500 times. By permuting sex labels, we break the association between sex-differential expression and sex-differential expression, while preserving other gene-level features. To quantitatively evaluate the significance of Twin Peaks in the null distribution, we used the three criteria laid out by Cheng and Kirkpatrick (2016) for classifying a curve as Twin Peaks. Namely, a 4^th^-degree polynomial must 1) be significant (*p* < 0.05) for the 4^th^-degree term by ANOVA, 2) have a negative coefficient for the Δ^4^ term, and 3) have three real roots. Any 4^th^-degree regression passing all three criteria is classified as Twin Peaks.

### Applying a new model and method for SDS-expression regression

In the section “**No evidence for SDS on eQTLs across all genes in multiple tissues**”, we described a revised method for testing a genome-wide relationship between SDS and Δ. For this analysis, we revised the inference model and data. We used the Non-Finnish European subset in the gnomAD V3 dataset^31^ to calculate between-sex *F*_ST_ and heterozygosity at each eQTL. This set of samples has an order of magnitude more individuals (average of 31,470 samples per site) compared to 1000 Genomes data (average of 2,300 samples per site when combining all ancestry groups) originally used in CK16. Additionally, we used the normalized TPM from the current GTEx v8 for expression level data, which includes many more samples (average of 236 male and 168 female samples per tissue) than the pilot (8 male samples and 13 female samples in the gonads)^28^. This version also provides sample-specific sex labels, enabling us to use additional tissues beyond gonads.

The equation we base our regression on is the model relating *F*_ST_ with *δ*^2^ shown in **Eq. 2**. Now, rather than performing a 4^th^-degree polynomial regression as in the analysis for “**Twin Peaks is a Statistical Artifact**”, we perform a weighted linear regression of *F*_ST_ to *δ*^2^ = 4*pq*Δ^2^, isolating *A* as the coefficient of regression. Note that by including 4*pq* in the independent variable, we allow information about heterozygosity to affect the SDS-expression relationship. We weighted each point of the regression by 1/*Var*(*expression*), where *Var*(*expression*) is the averaged variance in expression of each sex. This should decrease the leverage of points with large |Δ|. To get a gene-wide estimate of *F*_ST_, we used eQTLs from GTEx as mapped in the v8 study^31^. For each gene, we chose the single *cis*-eQTL (within 1Mbp of the gene’s midpoint) with lowest *p*-value for association with the gene and use that site for calculating *F*_ST_, *p*, and *q*. By using an eQTL selected this way, we tried to isolate the effect of selection on gene expression to the site presumed to be contributing most to expression changes in that gene.

To determine the significance of our *A* estimates, we generated 90% null acceptance regions by permuting sex labels. We permuted sex labels as in “**Twin Peaks sex-label permutation**” such that for each tissue, sex labels are permuted in GTEx expression samples and Δ is recalculated across all genes. Then, for each iteration *A* is recalculated by linear regression using the new Δ values (*F*_ST_ and 4*pq* remain unchanged). The 90% null acceptance region was obtained by the 5^th^ and 95^th^ percentiles of 1,000 *A* values calculated on permuted expression data.

